# Lethal Sudan virus infection in IFNAR^-/-^ mice reveals hallmarks of a cytokine storm

**DOI:** 10.64898/2026.03.30.715315

**Authors:** Michelle Gellhorn Serra, Cornelius Rohde, Lucie Sauerhering, Lars Meier, Lennart Kämper, Pauline Neubecker, Markus Eickmann, Alexandra Kupke, Stephan Becker, Anke-Dorothee Werner

## Abstract

Sudan virus (SUDV) is a member of the family *Filoviridae*, which comprises highly pathogenic viruses associated with unusually high case fatality rates. The development of medical countermeasures against filoviruses, including antivirals, vaccines, and therapeutic antibodies, requires preclinical evaluation in suitable animal models. C57BL/6J IFNAR^-/-^ mice, which lack the type I interferon (IFN-α/β) receptor, have been reported to be susceptible to filovirus infections, although their impaired innate immune response may represent a potential limitation of the model.

Here, we show that IFNAR^-/-^ mice constitute a suitable model for SUDV infection. Following infection, animals developed a clear clinical disease characterized by significant weight loss and pronounced changes in behaviour and appearance. Mice reached the predefined clinical endpoint 3–5 days post infection. *Post mortem* analysis of terminal samples revealed high viral loads and viral genome copies in all tested organs as well as in serum, indicating widespread systemic dissemination. Importantly, infection was associated with a marked increase in several key chemokines and cytokines linked to systemic inflammation, consistent with the development of a cytokine storm–like response.

Together, these findings demonstrate that SUDV infection in IFNAR^-/-^ mice induces systemic viral dissemination and a pronounced inflammatory response, supporting the suitability of this model for investigating filovirus pathogenesis and infection-associated immune dysregulation.

## Introduction

The Sudan virus (SUDV) belongs to the order *Mononegavirales* and is a member of the genus *Orthoebolavirus* within the *Filoviridae* family. Since SUDV was first reported in 1976, the virus has periodically emerged, causing nine additional outbreaks with case fatality rates ranging between 41 and 70% (1, 2). SUDV and other filoviruses are listed as priority pathogens by the WHO R&D Blue-print initiative as they pose a great threat to public health care systems because of their epidemic potential and insufficient medical countermeasures (3). Most filoviruses are therefore classified as biosafety level-4 (BSL-4) pathogens.

Wild-type (WT) rodents are not susceptible to filoviruses and show minimal or no symptoms. After adaptation through serial passaging in rodents, filoviruses adopt genomic mutations which are accompanied by the development of severe disease (4–6). As a result, these adapted viruses may not accurately reflect WT pathogenesis and vaccines, antibodies or small molecule antivirals developed against adapted viruses might not be optimal to counteract WT virus infection. Immunocompromised rodents therefore represent a useful tool that supports WT filovirus infection efficiently (7, 8, 4). Common rodent models include the interferon-α/β receptor knockout (IFNAR KO), the double interferon-α/β and γ receptor (IFNAGR) KO, interferon-γ receptor (IFNGR) KO, the cytoplasmic signal transducer and activator of transcription-1 protein (STAT-1) KO or the severe combined immunodeficiency (SCID) KO mice. Their inherently compromised immune responses represent limitations regarding infection and disease progression, as detailed information on the mounted immune response in successfully infected animals is missing for a number of mouse and virus strain combinations. Nevertheless, IFNAR^-/-^ mice, for instance, can efficiently mount humoral and cellular immune responses, and are therefore commonly used to evaluate different vaccine strategies, including those for Zika virus (9), Ebola virus (8, 10, 11) and Crimean-Congo haemorrhagic fever virus (12). For SUDV, several of such immunocompromised mouse models exist with different combinations of mice and virus strains (4, 7, 8, 13–17). The lethality of SUDV infection in these models varies and the majority of these studies focused on haematology and serum chemistry.

In the present study, we have characterized cytokine and chemokine responses in an IFNAR^-/-^mouse model of SUDV Boniface infection, which share similarities with other animal models and human infection.

## Methods

### Cells and viruses

SUDV isolate Boniface was propagated on Vero C1008 cells (clone E6, VeroE6, ATCC CRL-1586, African green monkey kidney cell), cultivated in Dulbecco’s modified Eagle medium (DMEM) supplemented with 3% fetal bovine serum, 1% L-Glutamine (200 mM), and 1% penicillin (50 units/ml) and streptomycin (50 mg/ml) (DMEM ++) at 37 °C and 5% CO_2_. Viral titers were calculated based on TCID_50_ and plaque assays (using VeroE6 cells). The final virus stock was sequenced and tested negative for contaminations such as mycoplasma or other virus infections (not shown).

All experiments with SUDV were performed in the high containment facility (BSL-4) of the Institute for Virology, University of Marburg, Germany, according to national and international regulations.

### Mouse experiments

Type I interferon receptor knock-out (IFNAR^-/-^) mice (18) with a C57BL/6J background were selected because of their reportedly high susceptibility for filovirus infections (4, 8) and were generously provided by U. Kalinke (Twincore, Hannover). Mice of both sexes were obtained from our in-house breeding colony. All experiments and protocols were approved by the regional authorities *Regierungspräsidium Gießen* AZ V54 –19 c 20 15 h 01 MR 20/7 Nr. G 64/2024), conducted according to the recommendations of Federation of European Laboratory Animal Science Associations and *Gesellschaft für Versuchstierkunde* (Society for Laboratory Animal Science, GV-SOLAS) and in compliance with the German animal welfare act and Directive 2010/63/EU.

Mice were housed as described previously (19, 12).

Three female (F) and three male mice (M), aged 6 months, were briefly anesthetized using isoflurane (CP-Pharma, Burgdorf, Germany) and infected intraperitoneally (i.p.) with approx. 1,000 plaque forming units (PFU) of SUDV in DMEM at a total volume of 200 µL. Mice were checked daily for changes in appearance, behaviour or body weight. Blood samples were drawn from the facial vein 3 days post infection (dpi) and immediately prior to euthanasia. Upon reaching the humane endpoints (clinical score of 10 or higher or score of 6 on two consecutive days; see S1 Table for more detailed information), the mice were euthanized by cervical dislocation under isoflurane.

Tissue samples from mesenteric lymph nodes, spleen, liver, kidney, lung, thymus, eye and brain as well as ovaries or testicles and accessory genital glands were collected and used for downstream *post mortem* analyses.

### Determination of viral loads and titers

Tissue samples were homogenized in 1 mL DMEM with ceramic and glass beads (Lysing Matrix H 2-mL tubes, MP Biomedicals) in a Mixer Mill MM 400 (Retsch, Germany) for 5 min. Afterwards, the homogenates were centrifuged for 5 min at 2,400 rpm and the supernatant was used for the isolation and quantification of viral RNA using qRT-PCR and the determination of viral titers in the different organs via TCID_50_ method.

#### qRT-PCR of serum and tissues

Isolation and quantification of viral RNA was performed as previously described (20). Briefly, viral RNA was isolated using the RNeasy Mini Kit (Qiagen) according to the manufacturer’s instructions and purified using the Quick-DNA/RNA viral MagBead kit (Zymo research) according to manufacturer’s instructions along with the fully automatic nucleic acid extraction system (Tecan). Viral loads were assessed using the RealStar® Filovirus Screen RT-PCR Kit 1.0 (altona Diagnostics) according to manufacturer’s instructions (including the following adaptation: volumes for both master mix and RNA were halved) on a qTOWER (Analytik Jena).

#### Virus titration by TCID_50_

To determine the levels of infectious virus in organ homogenates or serum samples, VeroE6 cells were seeded in 96-well plates (1x 10^4^ cells/well) and infected with 10-fold serial dilutions of supernatants from either organ homogenates or serum samples. At 6 dpi the cytopathic effect (CPE) was analyzed and TCID_50_/ml were calculated according to Spearman and Kerber (21). The limit of detection (LOD) for the organs was calculated by using the minimal detectable TCID₅₀/ml titer possible and relating it to the heaviest organ sample measured. This approach yields the lowest possible detectable value. The result was then extrapolated to 25 mg of tissue, providing the final LOD expressed as TCID₅₀ per 25 mg of tissue.

### Histopathological examination

The histological analysis was performed as described previously (22, 23). Briefly, collected organs were fixed in formalin and embedded in paraffin. Sections with a 4 µM thickness were cut with a microtome and stained with haematoxylin and eosin (H&E) for histopathological analysis. Furthermore, SUDV-specific RNA was stained by *in situ* hybridization (ISH) using the RNAscope^®^ 2.5 HD Assay—RED Kit (Cat. No. 322360) from Bio-Techne, with the probes V-SudanEbola-NP-sense (Cat No. 479281) and a custom designed probe V-SudanEbola-NP-O1-C1 (1846971-C1). Signal amplification was then performed using alkaline-phosphatase–labelled probes in combination with Fast Red substrate, which allowed signal detection. In parallel, a Negative Control Probe (Cat No. 310043) was used to control for background staining. Finally, the slides were counterstained with Gill’s Hematoxylin I and 0.02% ammonia water. Histological samples were evaluated independently and semi quantitatively by a trained scientist and a blinded veterinary pathologist. For liver tissue, the following parameters were assessed: necrotic foci, immune cell infiltration, steatosis, eosinophilic liver cells, and structural tissue damage. For the spleen following parameters were evaluated: white pulp atrophy (lymphoid depletion), congestion, focal single-cell necrosis, and infiltration by polymorphonuclear cells. Each parameter was scored using a 0–3 scale, with higher scores indicating greater severity.

### Multiplex ELISA – Luminex technology

To analyze inflammatory responses, final serum samples were screened for the release of cytokines and chemokines using the Bio-Plex Mouse Cytokine 23-Plex Assay by BioRad, as previously described (24). Briefly, magnetic beads, each containing a unique mixture of red and infrared fluorophores for identification and coupled to specific antibodies for analyte detection, were transferred to the assay plate and washed twice. Then, standards, samples, and controls were added and incubated for 30 min, followed by 25 µl detection antibody solution incubated for 30 min and 50 µl streptavidin-phycoerythrin (PE) for 10 min. Finally, the samples were resuspended in 125 µl assay buffer and mixed for 30 s. Data were acquired on a BioPlex 200 system (BioRad Laboratories). The incubation steps were performed on a shaker at 850 r.p.m. and room temperature (RT). Washing steps were performed after each incubation step on a Bio-Plex Pro Wash Station (BioRad Laboratories). The analyzed cytokines included IL-1α, IL-1β, IL-2, IL-3 IL-4, IL-5, IL-6, IL-9, IL-10, IL-12 (p40), IL-12 (p70), IL-13, IL-17A, eotaxin, IFN-γ, TNF-α, KC, RANTES, MIP-1α, MIP-1β, MCP-1, G-CSF and GM-CSF (see S3 Table). A five-parameter logistic regression was used to calculate the concentrations of the different analytes in the samples. The lower and upper limit of quantification were defined as the lowest and highest standards for each analyte, respectively. Serum samples of historical non-infected C57BL/6J IFNAR^-/-^ mice were used as controls. For statistical analysis, the data were log-transformed and analyzed using an unpaired *t*-test to compare, for each analyte, the values from SUDV-infected mice with those from uninfected controls, using Graphpad Prism 8.0.2 software.

### Next generation sequencing

Libraries for Illumina next generation sequencing of SUDV Boniface were prepared using the Twist Total Nucleic Acids Library Preparation EF Kit 2.0 (Twist Bioscience) for Viral Pathogen Detection and Characterization, as well as the Twist Target Enrichment Protocol (Rev. 2.0) with the Comprehensive Viral Research Panel. Libraries were sequenced on an Illumina iSeq 100 system, paired end reads 2 x 151 bp. Quality- and adapter trimming of raw reads, mapping and consensus sequence generation were performed using Geneious Prime 2022 build-in tools (BBduk, Geneious mapper, consensus sequence generation highest quality with cutoff 60%, respectively). Detected variants were visually validated in the alignment view. Detailed sequencing strategy and primers are available on request. Sequencing revealed three mutations (g605a (NP E150K), t5061c (VP40 L203P) and c12750t (L, silent mutation) compared to the reference sequence (FJ968794).

## Results

### Sudan virus infection causes a uniformly lethal disease in IFNAR^-/-^mice

3 female and 3 male IFNAR^-/-^ mice were infected intraperitoneally (i.p.) with approx. 1,000 PFU of SUDV Boniface isolate. Disease progression was monitored by measuring body weight and determining the overall appearance and behaviour daily for 14 days (Fig 1A). Each clinical symptom was assessed on a scale from 0 to 10 with 10 representing the most severe. The additive clinical score summarized the results of the different parameters analyzed daily (Fig 1B and S1 Fig). The mice were euthanized when they reached clinical score values of ≥10 or ≥6 on two consecutive days.

**Fig 1.**
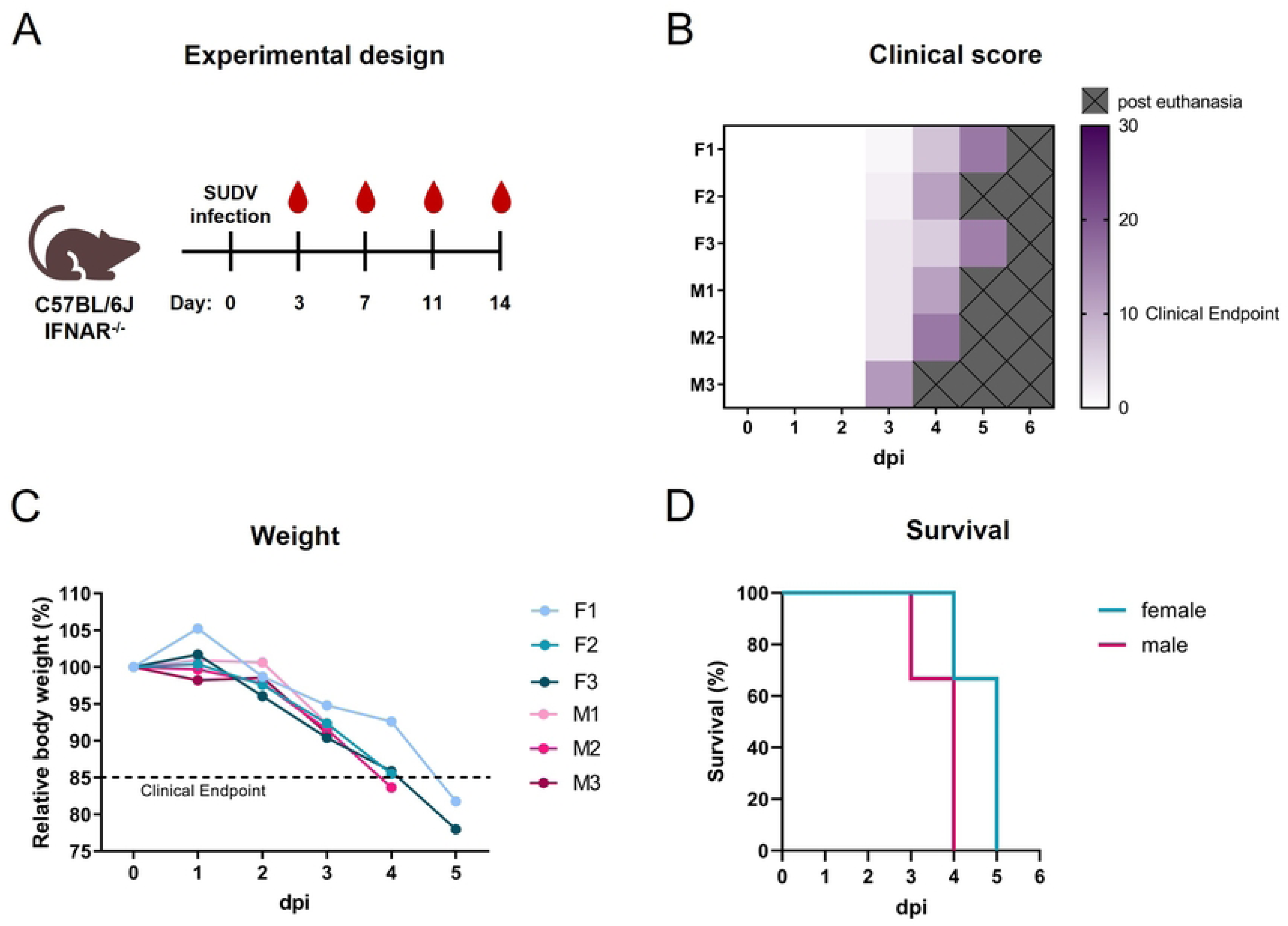
Clinical course of SUDV-infected IFNAR^-/-^ mice. The experimental approach is shown in **(A)**. Female (F) and male (M) IFNAR^-/-^ mice were infected with approx. 1,000 PFU SUDV on day 0 (n=6) and monitored daily. The mice were euthanized at humane clinical endpoint (score of ≥10 or ≥6 on two consecutive days). Blood samples were scheduled to be collected at day 3, 7 and 11, as well as on the day of euthanasia. The clinical score **(B)**, summarizes the results of the different parameters analyzed daily: body weight, appearance, and behaviour. Each parameter could be awarded a maximum of 10 points. Grey boxes indicate days when mice dropped out. The relative body weight compared to day 0 for each mouse is depicted in **(C)**. The dotted line marks the clinical endpoint. The Kaplan-Meier survival curve **(D)** depicts the percentage of surviving mice over time.

Mice began exhibiting signs of disease from day 3 onwards, with the most commonly observed symptoms being weight loss and reduced self-grooming behaviour (Fig 1B). Additionally, more severe symptoms included lethargy and reduced responsiveness (male mouse (M) 3) and haemorrhagic symptoms (female mouse (F) 3), bleeding from nostrils and oral cavity on day 5). Most mice reached the clinical endpoint due to marked weight loss of more than 15% (Fig 1C). Overall, the SUDV mouse model was characterized by 100% lethality according to the pre-defined humane end points within 5 dpi (Fig 1D).

### The Sudan virus infection results in widespread organ dissemination and high viral loads

During the course of the study, blood samples were collected from the facial vein on day 3 as well as on the day of euthanasia. Viral loads were assessed using qRT-PCR and TCID_50_ from serum and organ homogenates (Fig 2). For a better overview, the results for the qRT-PCR are depicted as 40-Ct values. Higher 40-Ct values indicate higher viral loads in the samples. As shown in Fig 2, substantial amounts of viral RNA were detected in all tested organs. The highest (40-Ct) values were found in the spleen (mean = 24.6), liver (mean = 22.9) and ovaries (mean = 23.61), while the lowest levels were found in the eyes (mean = 12.65) and brain samples (mean = 13.32). These values were relatively evenly distributed among the different organ samples, underscoring reproducibility and robustness of the model. qRT-PCR of serum samples showed similar values across all samples (mean = 17 on day 3), with the lowest results for F3 and M1 (dark green and light pink, respectively; Fig 2B). Missing values correspond to mice that have already dropped out.

**Fig 2.**
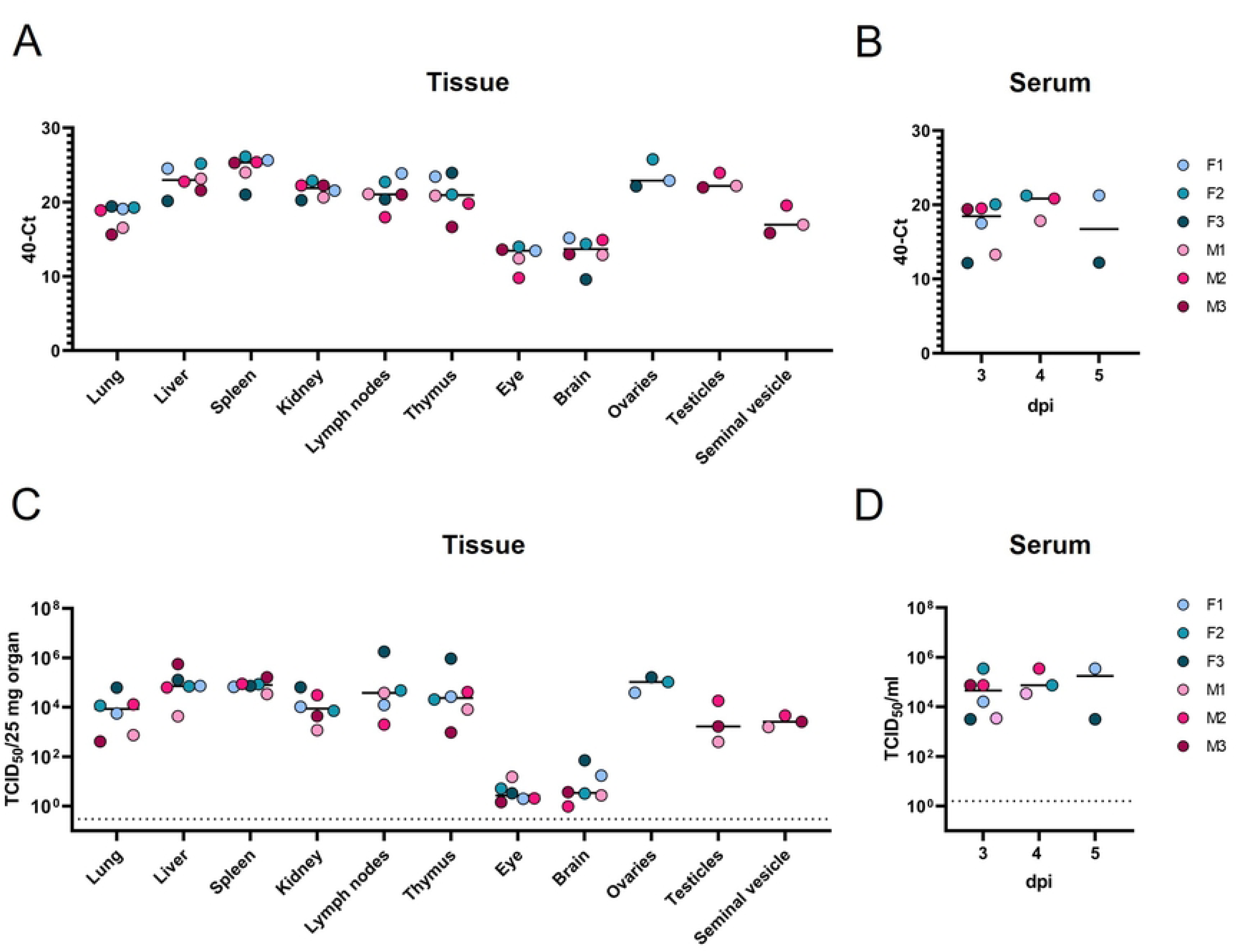
Detection of SUDV viral RNA and infectious titers in infected IFNAR^-/-^ mice. Blood samples were collected on day 3 and on the day of euthanasia, together with samples of different organs. The amounts of SUDV viral RNA were measured by RT-qPCR in the organs **(A)** and serum samples **(B)** of individual mice. The results are depicted as 40-Ct values for a better overview. Higher 40-Ct values indicate higher viral loads in the samples. Infectious SUDV titers were determined by TCID_50_ assay in organs **(C)** and serum samples **(D)** of individual mice. The titers present in the organs were normalised to their weight. Data are shown as mean with individual values. Dotted lines indicate the LOD (see the Methods section for details). **F:** female; **M:** male.

Infectious virus was also detected in all organs (Fig 2C) and in all serum samples (Fig 2D), confirming systemic infection. However, in the case of mouse M3, the titer for the lymph nodes was below the LOD due to an unspecific toxic effect on cells in the first dilution. The organs with the highest titers included liver (mean = 1,5×10^5^ TCID50/25 mg organ), spleen (mean = 8,6×10^4^ TCID50/25 mg organ), and ovaries (mean = 1×10^5^ TCID50/25 mg organ). On day 3, the average serum titer was 8.6 x 10⁴ TCID₅₀/ml. This increased further for surviving animals at 4 and 5 dpi (mean = 1.6 x 10⁵). Overall, the qRT-PCR results are consistent with the TCID_50_ values, indicating high viral RNA loads and infectious particles in all organs. This suggests that all tested organs supported productive infection.

Moreover, the harvested organs were analyzed histopathologically using H&E staining. Of the organs evaluated, the liver and spleen were the most affected and showed marked evidence of tissue damage. In the liver (Fig 3A-D), randomly distributed multifocal hepatocellular necrosis was evident (dotted lines), characterized by hepatocytes with hypereosinophilic cytoplasm (filled arrows, dark blue) and pyknotic or karyorrhectic nuclei (filled arrowheads, dark blue). In addition, variable and partially severe macrovesicular (and/or microvesicular) vacuolation of hepatocytes was noted (hollow arrows, dark blue), accompanied by mixed inflammatory-cell infiltration (hollow arrowheads). In the spleen (Fig 3E-H), the white pulp exhibited marked lymphoid depletion, (filled arrows, dark grey), while the red pulp contained foci with necrotic cells (filled arrowheads, dark grey) and patchy infiltration with mononuclear cells (sinus histiocytosis, hollow arrowheads, dark grey). Tissue damage was evaluated semi-quantitatively and scored from 0-3, with a score of 3 indicating greater severity. The highest scores for the liver and spleen were observed in mice F1 and F3, which survived the longest, whereas mouse M3, which dropped out on day 3, showed the lowest score for both organs. This pattern suggests that the severity of pathology correlates with the duration of infection, with mice that survived longer exhibiting more time-dependent progression of tissue damage.

**Fig 3.**
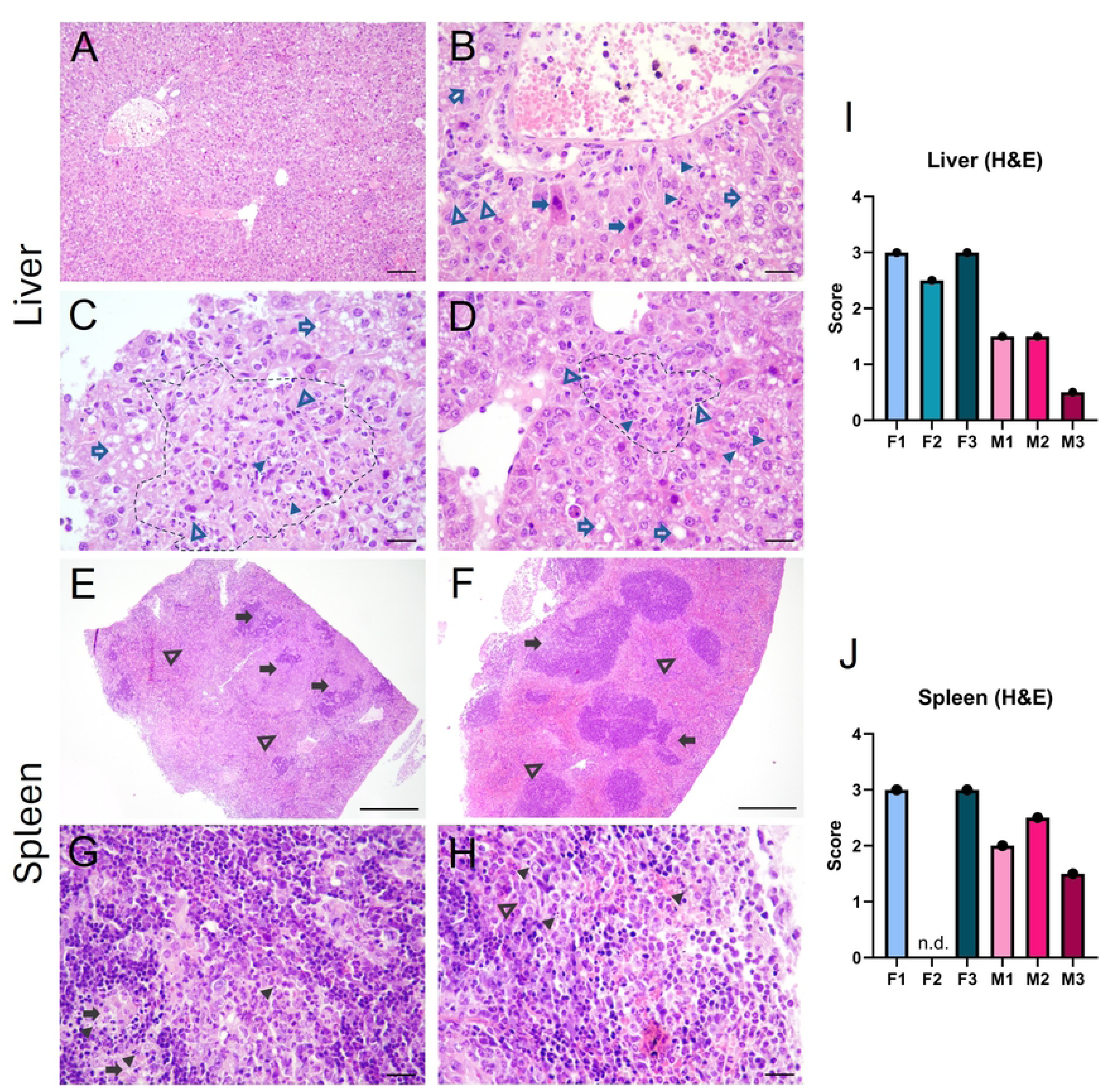
Pathology of SUDV-infected IFNAR^-/-^ mice. Organs were harvested after euthanization and pathologically examined *post mortem*. Exemplary images of liver **(A-D)** and spleen **(E-H)** stained with H&E are shown. In livers, multifocal liver necrosis, eosinophilic hepatocytes, lymphohistiocytic infiltrates and vacuolation were observed. In spleens, lymphoid depletion, necrotic cells as well as mononuclear infiltrates were observed. For each animal, scores for disease progression were determined (**I and J** for liver and spleen, respectively). Scale bars: A: 100 µm; B-D, G-H: 25 µm; E-F: 500 µm. **H&E:** hematoxylin and eosin; **n.d.:** no data; **F:** female; **M:** male.

The presence of genomic SUDV RNA in harvested organs was assessed via *in situ* hybridization using the V-SudanEbola-NP-sense probe, which hybridizes to antisense RNA. Red staining indicative of SUDV RNA was detected in multiple organs, with the most prominent staining observed in liver and spleen (Fig 4A-D). In these tissues, SUDV RNA appeared in small, scattered foci throughout the parenchyma, although the extent varied among individual mice. Notably, M1 and M2 exhibited strong staining in both liver and spleen, whereas M3 and F2 displayed only limited signal. In the brain, discrete foci positive for viral RNA were observed in the cortex, meninges and the choroid plexus (Fig 4E-F). Occasionally, signals for viral RNA were also detected in the lungs (Fig 4G-H), kidneys, visceral fat tissue, and testes. While overall detection of viral RNA levels was low, it was sufficient to induce severe tissue damage, likely driven by elevated and uncontrolled inflammatory responses. Interestingly, mice with the highest RNA signal received the lowest scores for tissue damage (S3 Fig and S2 Table). The inverse relationship between ISH RNA signal and tissue damage, combined with the observation that the most extensive lesions occurred in mice surviving the longest, may indicate that viral RNA decreases as tissue destruction progresses. Thus, heavily damaged tissues may no longer retain detectable levels of viral RNA.

**Fig 4.**
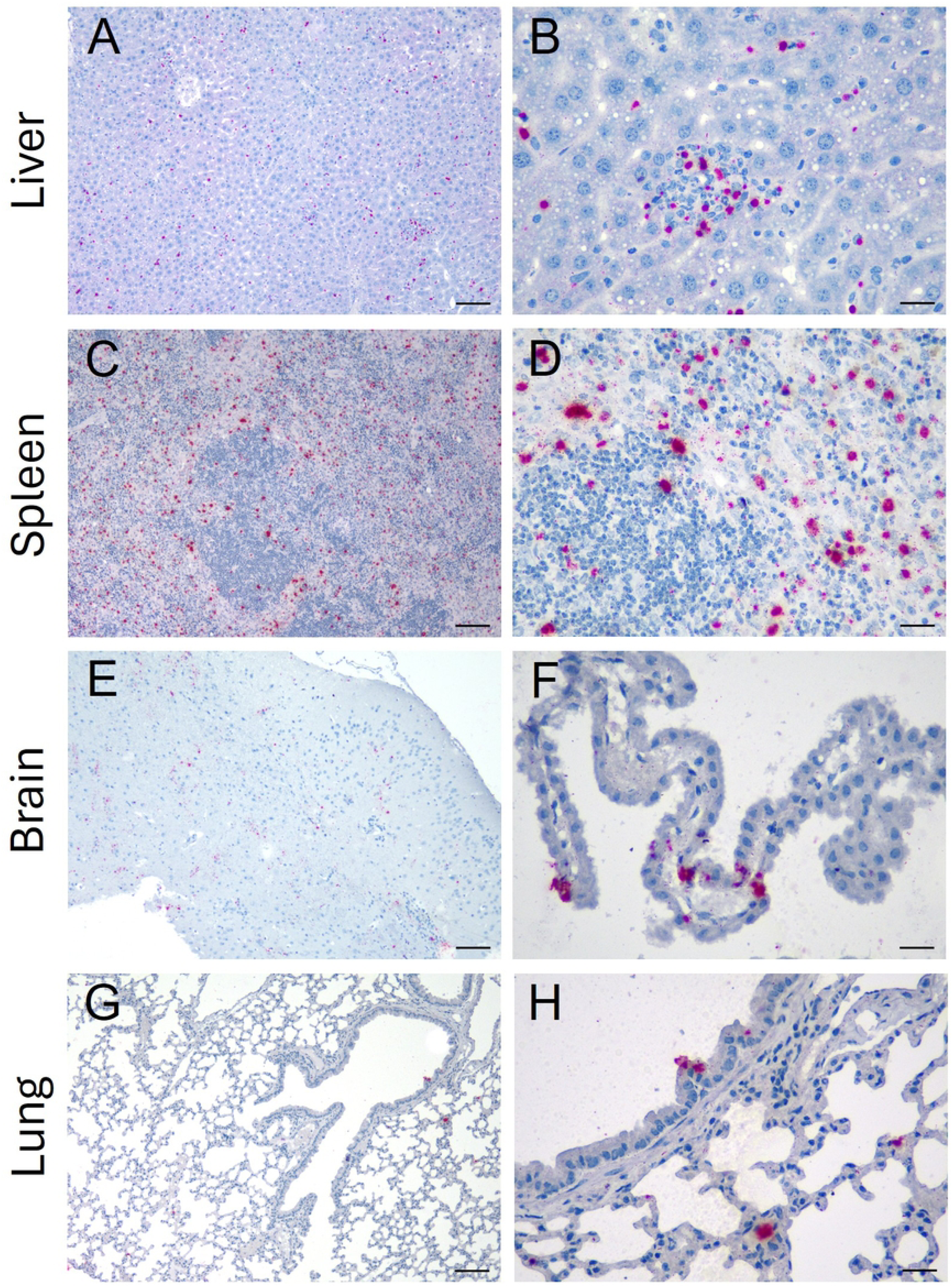
Detection of SUDV viral RNA in organs of IFNAR^-/-^ mice by *in situ* Hybridization. Organs were harvested after euthanization and pathologically examined *post mortem*. Exemplary images of liver **(A-B)**, spleen **(C-D)**, brain **(E-F)**, and lungs **(G-H)** stained for viral genomic RNA (V-SudanEbola-NP-sense) by ISH are shown (red staining). Scale bars: Left panel 100 µm; right panel 25 µm. **ISH:** *in situ* Hybridization.

Additionally, a custom-made probe was designed to detect SUDV Boniface but the results were comparable to the previously used probe (data not shown).

In summary, these results confirmed published data, showing IFNAR^-/-^ mice to be a suitable model for SUDV infections (4, 7, 8, 13–16).

### Sudan virus infection induces broadly elevated cytokine and chemokine responses

Fatal filovirus infections are characterized by immune suppression and dysregulated inflammatory responses (25, 26). To investigate whether this pattern is recapitulated in our model, we performed a chemokine/cytokine analysis and tested different pro-inflammatory cytokines, regulatory and anti-inflammatory cytokines, T-cell differentiation-associated cytokines, hematopoietic growth factors and chemokines for chemotaxis and recruitment, summarized in S3 Table (27).

As shown in Fig 5, all analytes in our panel were significantly upregulated in the SUDV-infected mice compared to our historical uninfected controls. Mouse F3 was not included in this analysis due to insufficient serum volume. Of note, male mice exhibited higher concentrations of several mediators than female mice in many instances, both under infected and control conditions. Moreover, mediators associated with different types of immune responses were significantly increased, including IFN-γ and IL-12 driving type I immunity; IL-4, IL-5, and IL-13 driving type II immunity; and IL-17a driving type III immunity. This simultaneous activation indicates a dysregulated immune response that can weaken or misdirect the type I immunity – which is crucial for clearing viral infections – ultimately impairing viral clearance and promoting pathological inflammation (28).

**Fig 5.**
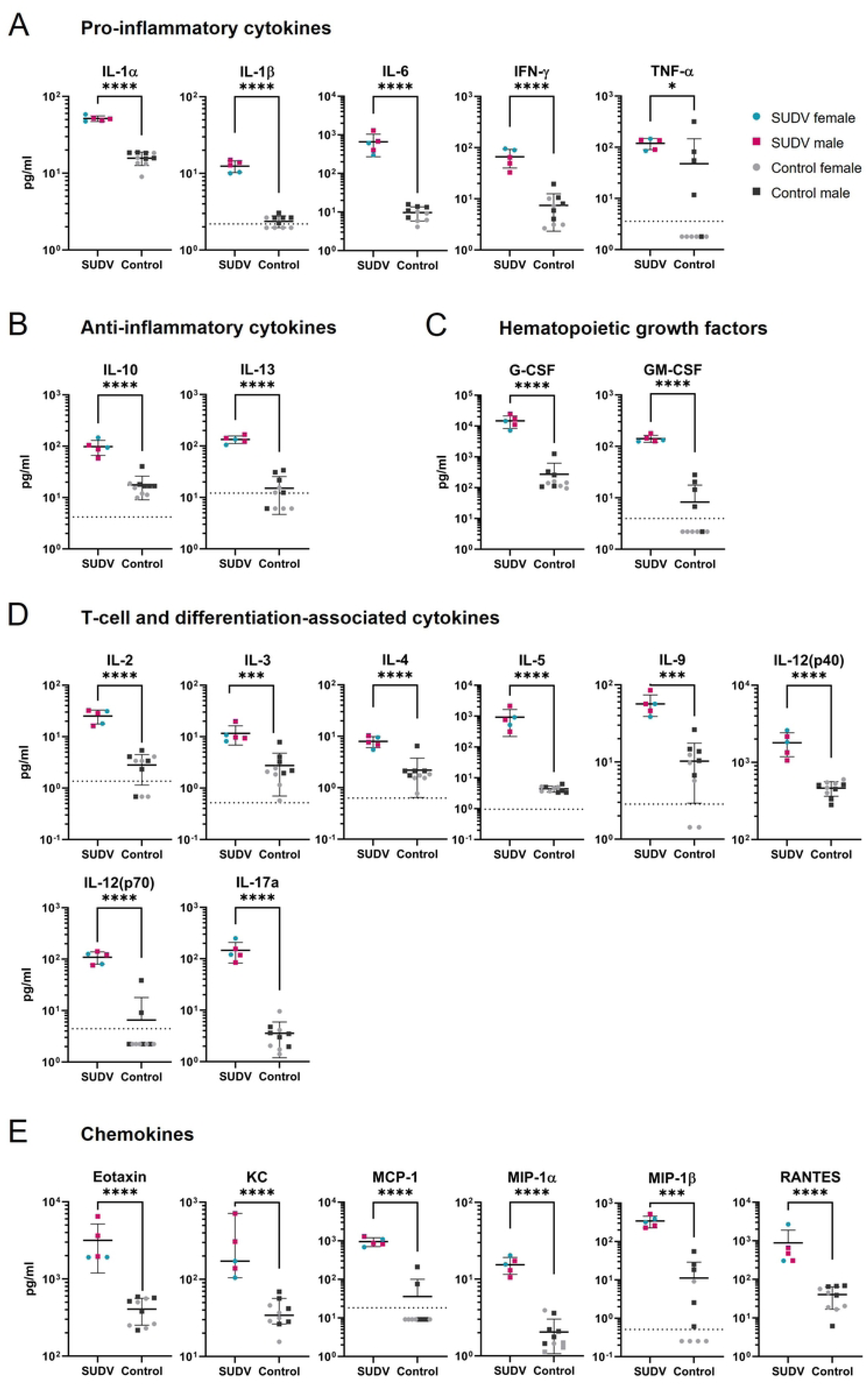
Systemic cytokine and chemokine response in SUDV-infected IFNAR^-/-^ mice. The presence of 23 mediators in the final serum samples of SUDV-infected mice and uninfected controls (n=10) was analyzed using multiplex ELISA based on the Luminex technology. Data are presented as mean ± SD, with individual values shown. Dotted lines indicate the lowest standard, defined as LLOQ. The data were log transformed for normal distribution prior to statistical analysis. The *p*-values are shown above each graph and refer to unpaired t-tests (see the Methods section for details). **ELISA:** enzyme-linked immunosorbent assay; **SD:** standard deviation; **LLOQ:** lower limit of quantification.

## Discussion

Here we present a uniformly lethal C57BL/6J IFNAR^-/-^ mice model for SUDV Boniface, where mice reached the clinical endpoint between day 3 to 5 after infection and were sacrificed with clinical scores of at least 10. The mice in this study were sacrificed earlier than in other studies due to stricter humane endpoints. Reported outcomes vary widely across models: Brannan et al. (2015) and Furuyama et al. (2016) found high survival in WT and IFNAR^-/-^ mice infected with SUDV Gulu or Boniface, whereas Froude et al. (2018), Comer et al. (2019), Escaffre et al. (2021), and Flaxman et al. (2024) reported uniformly lethal infections with varying disease onset and weight loss. Escudero-Pérez et al. (2019) observed ∼71% lethality in humanized mice, and Lever et al. (2012) reported non-lethal infection with ∼15% weight loss after airborne infection (8, 13–17, 7, 29).

Overall, SUDV Gulu and Boniface cause severe disease and high mortality, although outcomes vary depending on euthanasia criteria, mouse strain, and virus isolate. Only one study has directly compared the two SUDV strains, suggesting a potentially higher virulence of SUDV Boniface, although additional comparative studies are needed to confirm this observation (8). SUDV infection resulted in high viral RNA loads and infectious particles in serum and all analyzed tissues, indicating a systemic infection. The highest loads were detected in liver and spleen, followed by lymph nodes and thymus, consistent with previous reports of filovirus infection in humans showing prominent viral replication in secondary lymphoid organs, spleen, and liver (26). As has been reported for human patients infected with SUDV, fatal cases correlate with high viral RNA loads (30, 31), which has been used as a predictor of outcome. This is also reflected in our model, where we observed 100% lethality and high viral loads. Notably, viral RNA was also detected in reproductive organs (ovaries, testes, and accessory glands). As reports of viral persistence and sexual transmission have increased (32), these findings may be relevant but require further investigation. A mouse model for Ebola virus (EBOV) sexual transmission recently developed by Clancy and colleagues (33) may provide insights.

Histopathology revealed severe hepatosplenic damage. However, ISH staining for viral RNA was overall less extensive compared to EBOV-infected IFNAR^-/-^ mice (10). Similarly, Ellis and colleagues (34) found EBOV caused more widespread lesions and higher viral loads than SUDV in rhesus monkeys.

Variation in ISH staining among mice potentially reflects differences in euthanasia timing as mice euthanized later (M1 and M2, day 4) showed the highest loads, while one mouse euthanized earlier (M3, day 3) showed the lowest levels.

Slight sex-dependent differences were observed. For instance, male mice reached the humane endpoint on average one day earlier than female mice, and showed, in some instances, higher serum cytokine and chemokine concentrations. However, due to the limited number of animals, no definitive conclusions can be drawn. Further studies with larger cohorts will be required to clarify these observations. Nevertheless, sex-specific differences in basal cytokine and chemokine levels, as well as in responses to noxious stimuli including infection, injury, or inflammation, have been described previously (35–38). Male mice often exhibit stronger pro-inflammatory cytokine responses (e.g. IL-6, TNF-α) to infection or injury, which are associated with more pronounced sickness behavior. Conversely, female mice tend to display higher levels of specific chemokines that enhance leukocyte recruitment and immune cell reactivity, often yielding more effective, but potentially more damaging, immune responses.

Fatal filovirus infections of humans are characterized by immune suppression, uncontrolled inflammation, and weak humoral responses, whereas survivors show early and moderate immune activation with detectable IgG and IgM (39, 25, 26). Infected macrophages and monocytes release cytokines and chemokines such as TNF-α, IL-1β, IL-6, IL-8, IL-10, MCP-1, MIP-1α, MIP-1β, and IP-10, driving a dysregulated immune response, termed “cytokine storm” that can cause systemic damage and multi-organ failure (39–41). Fatal cases also show suppressed IFN-α, elevated pro-inflammatory mediators, and loss of CD4⁺/CD8⁺ T cells with reduced T-cell cytokines 13 (42, 43). Especially elevated levels of IL-6, IL-10 and MIP-1β have been specifically shown by Hutchinson and Rollin in 2007 and McElroy et al. for patients infected with SUDV in 2000/2001, using a similar Luminex technology as in the present study (44, 31). Consistent with these findings, the SUDV-infected mice of this study exhibited high serum cytokine and chemokine levels and pronounced hepatosplenic pathology, indicative of a cytokine storm not previously described in such detail for SUDV in mice (27).

Escudero-Pérez et al. compared EBOV and Reston virus (RESTV) infection in humanized mice using Luminex cytokine profiling. Serum data at 3 and 6 dpi are most comparable to our study. Relative to their EBOV data, the IFNAR^-/-^ mice in the present study showed similar IFN-γ and IL-10 levels, higher IL-6, RANTES, and MCP-1, but lower MIP-1α (S4 Table). In RESTV-infected mice, cytokine levels were similar or slightly lower than in EBOV infections, with markedly higher levels in non-survivors. EBOV cytokines generally rose over time, while RESTV levels stabilized or declined. If technical and strain differences between the two studies are neglected, cytokine expression ranked roughly RESTV < SUDV < EBOV. Since RESTV and EBOV mice were euthanized later, cytokine levels may converge near terminal stages (17). Luminex analysis of BALB/c mice infected with mouse-adapted Marburg virus (MARV) Angola showed significantly higher cytokine and chemokine levels than the symptomless wild-type strain at 6 dpi (45), though the magnitude of induction differed markedly from our SUDV results (S4 Table).

In Cynomolgus macaques, SUDV infection triggered strong cytokine and chemokine responses, broadly consistent with those observed in our mouse model (46). Notably, TNF-α, IFN-γ, and IL-2 - unchanged in human SUDV cases - were nonetheless elevated in both macaque studies and in our data, though to varying degrees. A second non-human primate study reported similar overall trends, with only modest differences in magnitude across individual analytes (47).

Luminex analysis of SUDV (Gulu)-infected human patients showed significantly elevated IL-1β, IL-6, IL-10, IP-10, MIP-1β, and RANTES, whereas TNF-α, IL-2, and IFN-γ were not upregulated compared to healthy controls (44). This contrasts with our SUDV study and EBOV reports (42, 25), although TNF-α was the least affected parameter in our study (Fig 5). Another study of the 2000–2001 Uganda outbreak found increased IL-1α, IL-6, MCP-1, and MIP-1α in fatal cases, consistent with our results, while IL-8 (KC homolog) rose only at late infection stages (31). Elevated M-CSF, MIP-1α, and IP-10 were linked to hemorrhages, whereas TNF-α was not significant and IL-2 and IFN-γ were excluded from analysis (31).

S4 Table summarizes cytokine and chemokine regulation across species.

The increase of TNF-α, IFN-γ, and IL-2 in macaques and our mouse model, but not in humans, raises questions about their role in SUDV pathogenesis, especially given their known upregulation in EBOV infection. However, controlled animal studies allow for precise timing of sample collection, whereas human samples are typically obtained at variable and often late stages of disease. These differences in sampling time may obscure smaller but biologically meaningful cytokine changes in human infections, especially because TNF-α is mainly present in the acute phase of infection and decays quickly (42).

IFNAR^-/-^ and similar KO mice are often a subject of critique as their immune system is different from WT mice which raises the questions whether KO mice are indeed suitable models to study filovirus infection. A feasible attempt to avoid both KO mice and mouse-adapted viruses is the treatment of (WT) mice with IFN antibodies prior to infection to enable/enhance susceptibility (4). However, the fact that filovirus infections result in a similar chemokine/cytokine profile independent on the mouse strain, and that these profiles are comparable between KO and WT mice and even comparable to NHPs and humans, strengthens the relevance and comparability of IFNAR^-/-^ mouse models to wild-type systems in the study of filovirus-induced inflammatory responses.

In summary, we present new data supporting the IFNAR^-/-^ mouse as a suitable model for SUDV Boniface infection. This model produces a uniformly lethal disease with hallmarks typical of filovirus infection. Elevated cytokine and chemokine levels indicate a cytokine storm, closely resembling responses observed in human and non-human primate infections supporting the model’s value for future vaccine and antiviral efficacy studies.

## Abbreviations

BSL-4: biosafety level-4;
Dpi: days post infection;
EBOV: Ebola virus;
ELISA: enzyme-linked immunosorbent assay;
G-CSF: granulocyte-colony stimulating factor;
GM-CSF: granulocyte-macrophage colony-stimulating factor;
IFN-γ: interferon gamma;
IFNAR-/-: interferon receptor alpha/beta knockout mice;
IL: interleukin;
i.p.: intraperitoneally;
KC: keratinocyte-derived chemokine;
LLOQ: lower limit of quantification;
LOD: limit of detection;
MARV: Marburg virus;
MCP-1: monocyte chemotactic protein-1;
MIP: macrophage inflammatory protein;
PFU: plaque forming units;
RANTES: Regulated upon Activation, Normal T cell Expressed and Secreted;
RESTV: Reston virus;
SUDV: Sudan virus;
TNF-α: tumor necrosis factor alpha.

## Acknowledgements

The authors would like to thank Gotthard Ludwig and Sebastian Schmidt as well as Astrid Herwig and Katharina Kowalski for excellent technical assistance. This project is funded by the State of Hesse, LOEWE Center "DRUID" (project A1), Flexfund "Establishment of a Sudan Ebola virus mouse model”, “UNISCIENTIA STIFTUNG VADUZ” and by the DZIF (Deutsches Zentrum für Infektionsforschung) TTU Emerging Infections (FKZ 8033801809). This project was also co-funded by the European Vaccines Hub for Pandemic Readiness, which is funded by the European Union. Views and opinions expressed are, however, those of the authors only and do not necessarily reflect those of the European Union or European Health and Digital Executive Agency (HaDEA). Neither the European Union nor the granting authority can be held responsible for them.

## Author contributions

Conceptualization: MGS, AW, CR, SB; Data curation: MGS, AW, CR LS, LK, LM, PN, ME, AK, ME; Funding acquisition: AW, SB; Project administration: AW, SB; Visualization: MGS, AW; Writing – original draft: MGS, AW, SB; Writing – review & editing: all authors

## Inclusion and diversity

We support inclusive, diverse, and equitable conduct of research.

